# Connecting the dots between root cross-section images and modelling tools to create a high resolution root system hydraulic maps in *Zea mays*

**DOI:** 10.1101/2020.12.15.422825

**Authors:** Adrien Heymans, Valentin Couvreur, Guillaume Lobet

**Affiliations:** Earth and Life Institute, UCLouvain, Louvain-la-Neuve, BE; Agrosphere Institute, Forschungszentrum Juelich, Juelich, DE

**Keywords:** Root anatomy, hydraulic conductivity, hydrophobic barriers, GRANAR, MECHA

## Abstract

Root hydraulic properties play a central role in the global water cycle, agricultural systems productivity, and ecosystem survival as they impact the global canopy water supply. However, the available experimental methods to quantify root hydraulic conductivities, such as the root pressure probing, are particularly challenging and their applicability on thin roots and small root segments is limited. There is a gap in methods enabling easy estimations of root hydraulic conductivities across a diversity of root types and at high resolution along root axes.

In this case study, we analysed *Zea mays* (maize) plants of the var. *B73* that were grown in pots for 14 days. Root cross-section data were used to extract anatomical measurements. We used the Generator of Root Anatomy in R (GRANAR) model to generate root anatomical networks from anatomical features. Then we used the Model of Explicit Cross-section Hydraulic Anatomy (MECHA) to compute an estimation of the root axial and radial hydraulic conductivities (*k_x_* and *k_r_*, respectively), based on the generated anatomical networks and cell hydraulic properties from the literature.

The root hydraulic conductivity maps obtained from the root cross-sections suggest significant functional variations along and between different root types. Predicted variations of *k_r_* along the root axis were strongly dependent on the maturation stage of hydrophobic barriers. The same was also true for the maturation rates of the metaxylem. The different anatomical features, as well as their evolution along the root type add significant variation to the *k_r_* estimation in between root type and along the root axe.

Under the prism of root types, anatomy, and hydrophobic barriers, our results highlight the diversity of root radial and axial hydraulic conductivities, which may be veiled under low-resolution measurements of the root system hydraulic conductivity. While predictions of our root hydraulic maps match the range and trend of measurements reported in the literature, future studies could focus on the quantitative validation of hydraulic maps. From now on, a novel method, which turns root cross-section images into hydraulic maps will offer an inexpensive and easily applicable investigation tool for root hydraulics, in parallel to root pressure probing experiments.

**One-Sentence summary:** The use of cross-section images and modelling tools to generate a map the axial and radial hydraulic conductivity along different root types for the maize cultivar B73.

## Introduction

Root hydraulics properties are one of the major functional plant properties influencing the root water uptake dynamics. Indeed, the radial hydraulic conductivity (*k_r_*) is a key component of the water absorption and the axial hydraulic conductance (*k_x_*) defines the water transport along the root (Leitner et al., 2014). Changes in the local root hydraulic properties, at the cell and organ scale, are known to have global repercussions on the root hydraulic behavior (Tardieu et al., 2018; Meunier et al., 2020) and are considered as important breeding targets to create drought resilient varieties (Maurel and Nacry, 2020). The quantification of root hydraulic conductivity along the roots is therefore needed to have a quantitative understanding of the root water uptake dynamics.

The root radial conductivity is influenced by different factors. For instance, root anatomical features define the baseline for radial water flow (Steudle, 2000; Heymans et al., 2019). The modulation of aquaporin can modulate that baseline value by affecting the cell membrane permeability on the short term (Parent et al., 2009). On the long term, the development of hydrophobic barriers (Enstone et al., 2002) and the conductivity of plasmodesmata (Couvreur et al., 2018) have also a crucial impact. On the other hand, the axial root conductance is a function of the xylem vessel area, maturation and number (Martre et al., 2001).

The quantification of radial hydraulic properties is challenging due to the complexity of the experimental procedures. It is even more complicated to assess it at different locations along the root axis and on different root types. The most direct way to estimate root radial conductivity is with roots which grow in soil-less environments using a root pressure probe (Frensch and Steudle, 1989). Other experimental techniques employed a pressure chamber to measure water flow that were successively cut into smaller parts (Zwieniecki et al., 2002), or employed the high pressure flow meter device (Tyree et al., 1994). Recently, virtual quantification of radial hydraulic properties was enabled with models such as the Model of Explicit Cross-section Hydraulic Anatomy (MECHA) (Couvreur et al., 2018). An intermediate technique uses inverse modeling method with the root architecture model of Doussan et al. (1998) and high resolution images of root water uptake (Zarebanadkouki et al., 2016). The estimation of axial hydraulic properties is easier than the radial ones since it can be calculated from Hagen-Poiseuille's equation with only a root cross-section image (Frensch and Steudle, 1989).

Since 1998, when Doussan et al. (1998) made a modelling approach to map the root hydraulic conductance on two *Zea mays* (maize) root types, little effort, to our knowledge, has been made to reproduce or to improve the spatial distribution of radial root hydraulic conductivity and axial root hydraulic conductance in maize.. However many studies that used functional-structural root model to simulate water uptake use the hydraulic conductivity that have been estimated by Doussan et al. (1998), such as in R-SWMS (Javaux et al., 2008), OpenSimRoot (Postma et al., 2017) or MARSHAL (Meunier et al., 2020). Although those estimations were groundbreaking for the community at the time, we now need to be able to quantify root hydraulic conductivities that directly match the data that we want to assess. Therefore including the effect of root anatomical changes and taking into account cell hydraulic properties would improve the accuracy and prediction of root water uptake models.

Here, we present a procedure to generate a high resolution hydraulic conductivity map from experimental data using recent modeling tools. With free hand cross sections and fluorescent microscopy, we were able to extract easily anatomical features that can be used to run the Generator of Root Anatomy in R (GRANAR) (Heymans et al., 2019). Then, using the generated anatomical networks with MECHA (Couvreur et al., 2018), we estimated the *k_r_* and *k_x_* along the root axis of each maize root type. This model's coupling creates a way to generate a root hydraulic conductivity map that takes into account the impact of the anatomy and the cell hydraulic properties. The method that we developed here does not rely on expensive equipment. It can be easily reproduced for other genotypes and different environmental constraints.

## Material and methods

Five *Zea mays* (maize cultivar B73) plants were grown in pots for 14 days. The pot dimensions were 12 cm diameter, 25 cm deep and filled with sieved potting soil. The soil was at field capacity when the germinated seeds were planted and never re-watered afterwards. The germination of the seed occurred in a petri dish maintained vertically in dark condition between two wet filter papers. From the fifteen seeds that were put under germination, five were selected based on the length of the tap root (0.5 to 1 cm long) in order to have an homogenous germination rate. Each seed was planted in a different pot. All plants grew in a greenhouse with the environmental settings of the greenhouse set to 60 % for the relative humidity and a temperature of 25°C (+− 3°C).

The root systems were excavated and washed at the end of the experiment (after 14 days). The root systems were scanned and selected root samples were conserved in a Formaldehyde Alcohol Acetic Acid solution (Ruzin and Others, 1999). The roots were stained with berberine for one hour and post stained with aniline blue for 30 minutes before making free-hand cross-sections (Brundrett et al., 1988). Three or more roots per type were cut at every 5 cm or less to map anatomical features along the root segments. Cross section images were acquired with fluorescent microscope SM-LUX and the pictures were taken using a Leica dfc320. The images were analysed with the ImageJ software. The anatomical features that we measured are listed in the table 1.

**Table 1:** List of the measured anatomical features acquired on the root cross section images that have been used to get the GRANAR parameters.

| Measured anatomical features | GRANAR parameters |
| --- | --- |
| epidermis cell width [$\mu\text{m}$] | epidermis *cell_diameter* |
| exodermis cell width [$\mu\text{m}$] | exodermis *cell_diameter* |
| cortex width [$\mu\text{m}$] | cortex *cell_diameter* |
| number of cortex cell layer [-] | cortex *n_layers* |
| endodermis cell width [$\mu\text{m}$] | endodermis *cell_diameter* |
| pericycle cell width [$\mu\text{m}$] | pericycle *cell_diameter* |
| stele diameter [$\mu\text{m}$] | stele *layer_diameter* |
| stele cell diameter [$\mu\text{m}$] | stele *cell_diameter* |
| metaxylem cell area [$\mu\text{m}^2$] | xylem *max_size* |
| number of metaxylem vessels | xylem *n_files* |
| number of protoxylem vessels | xylem *ratio* |

To identify the type of hydrophobic barriers that were encountered on the cross-section images, we used the berberine-aniline blue fluorescent staining procedure for suberin, lignin, and callose in plant tissue (Brundrett et al., 1988). This procedure for visualizing exo - and endodermal Casparian strips works also to identify the lignification of the xylem cell walls. Xylem vessels with fully lignified cell walls were considered as mature xylem elements.

The root type selected for this analysis are the tap root, the basal root (embryonic root), the shoot born root on the first node and two types of lateral roots, the short ones (short ones) and long ones (longer than 5 cm with second order lateral roots on it) (Passot et al., 2018). The choice of two classes of lateral root instead of three is due to experimental constraints. We had to base the classification on root length instead of root growth rate. The threshold that we set is evaluating the difference between the long later root classified as type A in the Passot et al. (2018) study and the other two lateral types (B and C) that have a slowing growth rate.

We modelled the evolution of anatomical features along the root axis and for different root types using linear models. The models were used to estimate the different GRANAR input parameters along the root axes. However, if the explanatory variable showed a p-value > 0.05, the average value of the anatomical features along the root axis was taken instead of the value predicted by the linear model. The generated anatomies were then used to estimate the kr and kx on each point along the selected spatial resolution for each root type using MECHA.

The statistical analysis was conducted using R (R Core Team, 2018). The R package that was used for the data analysis was “tidyverse” (Wickham et al., 2019).

### Description of MECHA Hydraulic Parameters

The simulation framework MECHA (Couvreur et al., 2018) can estimate root radial conductivities from the root anatomy generated with GRANAR and from the subcellular scale hydraulic properties of walls, membranes, and plasmodesmata. The cell wall hydraulic conductivity was set to 2.8 10^−9^ m^2^s^−1^MPa^−1^, as measured by Zhu and Steudle (1991) in maize. Lignified and suberized wall segments in the endodermis and exodermis were considered hydrophobic and attributed null hydraulic conductivities. The protoplast permeability (*L_pc_*, 7.7 10^−7^ m s^−1^MPa^−1^) measured by Ehlert et al. (2009) was partitioned into its three components: the plasma membrane intrinsic hydraulic conductivity (*k_m_*), the contribution of aquaporins to the plasma membrane hydraulic conductivity (*k_AQP_*), and the conductance of plasmodesmata per unit membrane surface (*K_PD_*). The latter parameter was estimated as 2.4 10^−7^ m s^−1^MPa^−1^ (Couvreur et al., 2018), based on plasmodesmata frequency data from Ma and Peterson (2001), and the plasmodesmata conductance estimated by Bret-Harte and Silk (1994). By blocking aquaporins with an acid-load treatment, Ehlert et al. (2009) measured a *k_AQP_* of 5.0 10^−7^ m s^−1^MPa^−1^. The remaining value of km after subtraction of *k_AQP_* and *K_PD_* from *L_pc_* was 0.3 10^−7^ m s^−1^MPa^−1^. Each value of km, *k_AQP_*, *k_PD_*, and *L_pc_* was set uniform across tissue types. For details on the computation of *k_r_*, see Couvreur et al. (2018).

The root axial hydraulic conductance was estimated using the Hagen-Poiseuille equations.

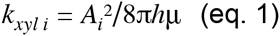

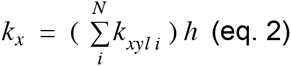

Where *A* is the cell area of one xylem vessel, *h* is the cell length and *μ* is the viscosity of the xylem sap. Xylem sap being essentially water, the viscosity constant was assumed to be equal to the one of the water.

As the root hydraulic conductivities obtained in this study are compared, among other studies, with the ones estimated in Doussan et al. (1998), we added an assumption to the data provided from that study. This hypothesis is that the lateral roots have an average growing rate of one centimeter per day (Passot et al., 2018).

The details about the GRANAR-MECHA coupling is available in an online Jupyter NoteBook (https://mybinder.org/v2/gh/HeymansAdrien/GranarMecha/main). The complete procedure can be run online or locally after downloading the related gitHub repository (https://github.com/HeymansAdrien/GranarMecha doi: 10.5281/zenodo.4316762). This complementary open-source ressources helps the users to change anatomical features and change cell hydraulic properties to personalise the exercise at will. The outputs of each generated root cross-section can be visualized through different figures that show the proportion of the water fluxes in each compartiment (apoplastic and symplastic fluxes).

The whole script that was used to compute the root hydraulic maps from the root anatomical measurement is presented as a Rmarkdown script stored in a GitHub repository (https://github.com/granar/B73_HydraulicMap doi: 10.5281/zenodo.4320861). In the same repository are stored all input and output data of this study.

## Results

To create hydraulic conductivity maps along the different maize root types taking into account the evolution of the anatomical features, we needed to capture anatomical descriptors that are ready-to-use for downstream computational models. Anatomical features change along the root axis, as the root is narrower and less mature at the tip than at its basal position. Across root types, anatomies may also differ. With the gathered root cross section images, we were able to extract the root anatomical features and place those features along the root axes. Most of the root anatomical features that we computed follow a linear regression when they are plotted against the distance to the tip (Figure 1, Suppl. Fig 1).

**Figure 1:**
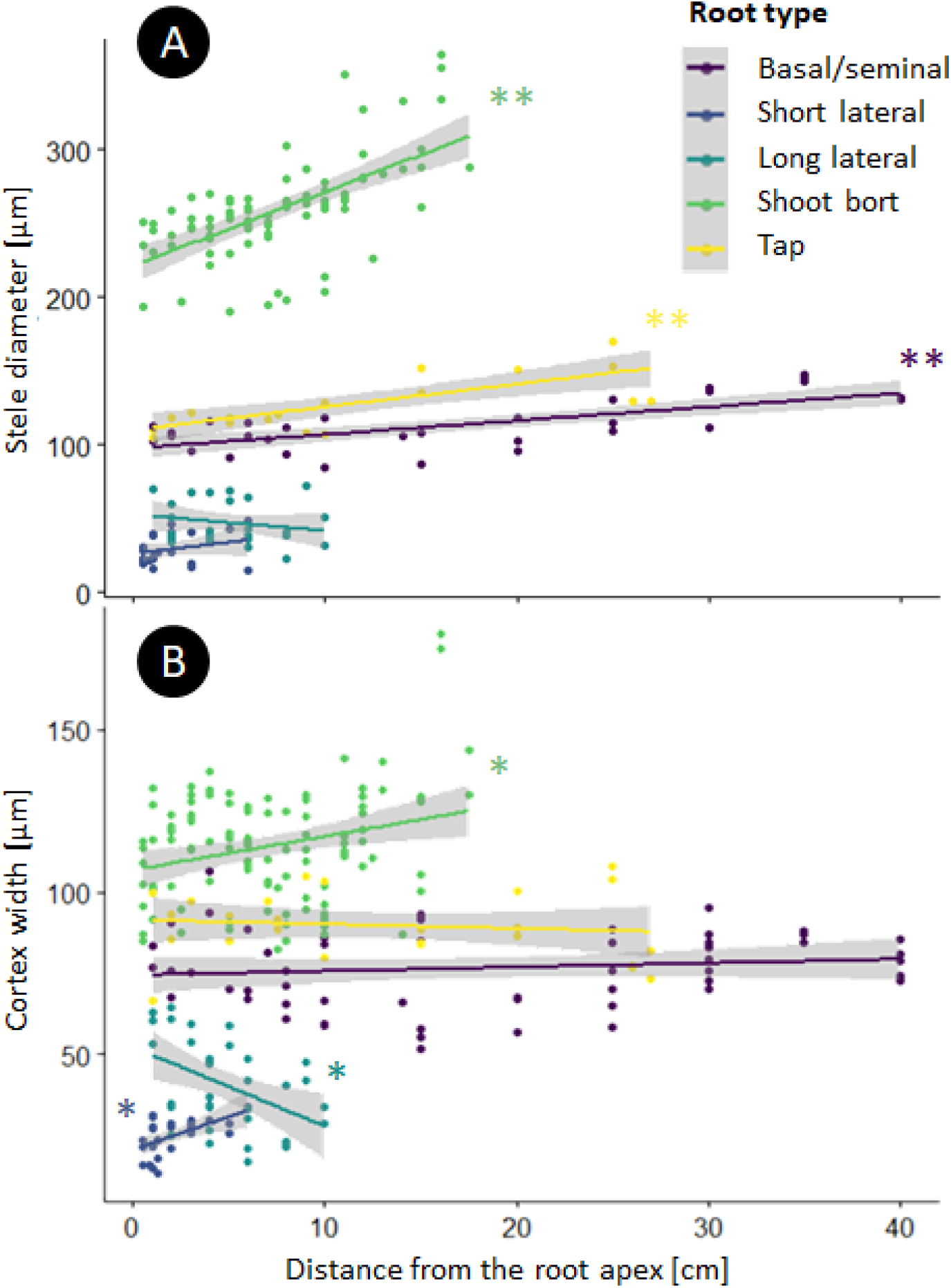
Evolution of the stele diameter (A) and the cortex width (B) along the root axe for the different root types. **: P <0.001; *: P< 0.01; ‘’: P > 0.05

The stele of the root axes (tap-, basal-, and shoot born-root) narrows close to the tip. As the stele area shrinks, the number of xylem vessels are also reduced. The correlation between the stele and xylem areas is strong (0.899) but it is not suitable to do a linear regression. However when we look at the Napierian logarithm of those areas (Yang et al., 2019), the linearity of this relationship is strong (R^2^: 0.9913, fig. 2). Thanks to the strong relationship between those anatomical features, we used it into the model parametrization procedure instead of using directly the xylem size data of the anatomical features previously measured.

**Figure 2:**
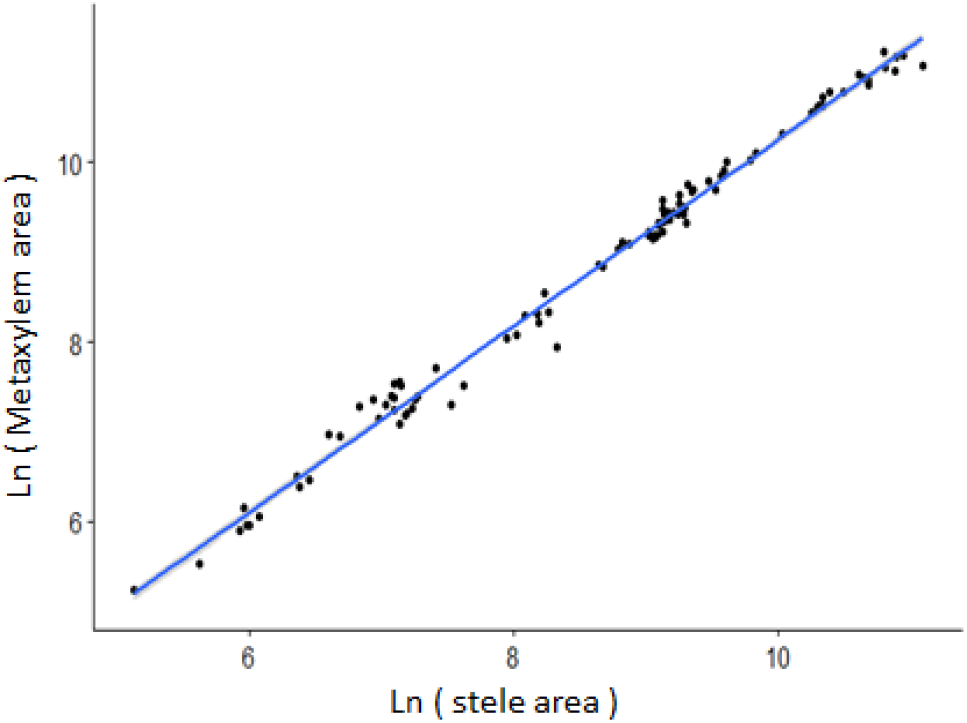
Allometric relationship between the metaxylem area and the stele area. Both area are expressed in mm^2^

For each of the GRANAR model parameters we have a simple function that depends on the root type and the distance from its apex (Supplemental Table 1). With that information, we were able to build average root cross sections along each root type, at any longitudinal position (fig. 3).

**Figure 3:**
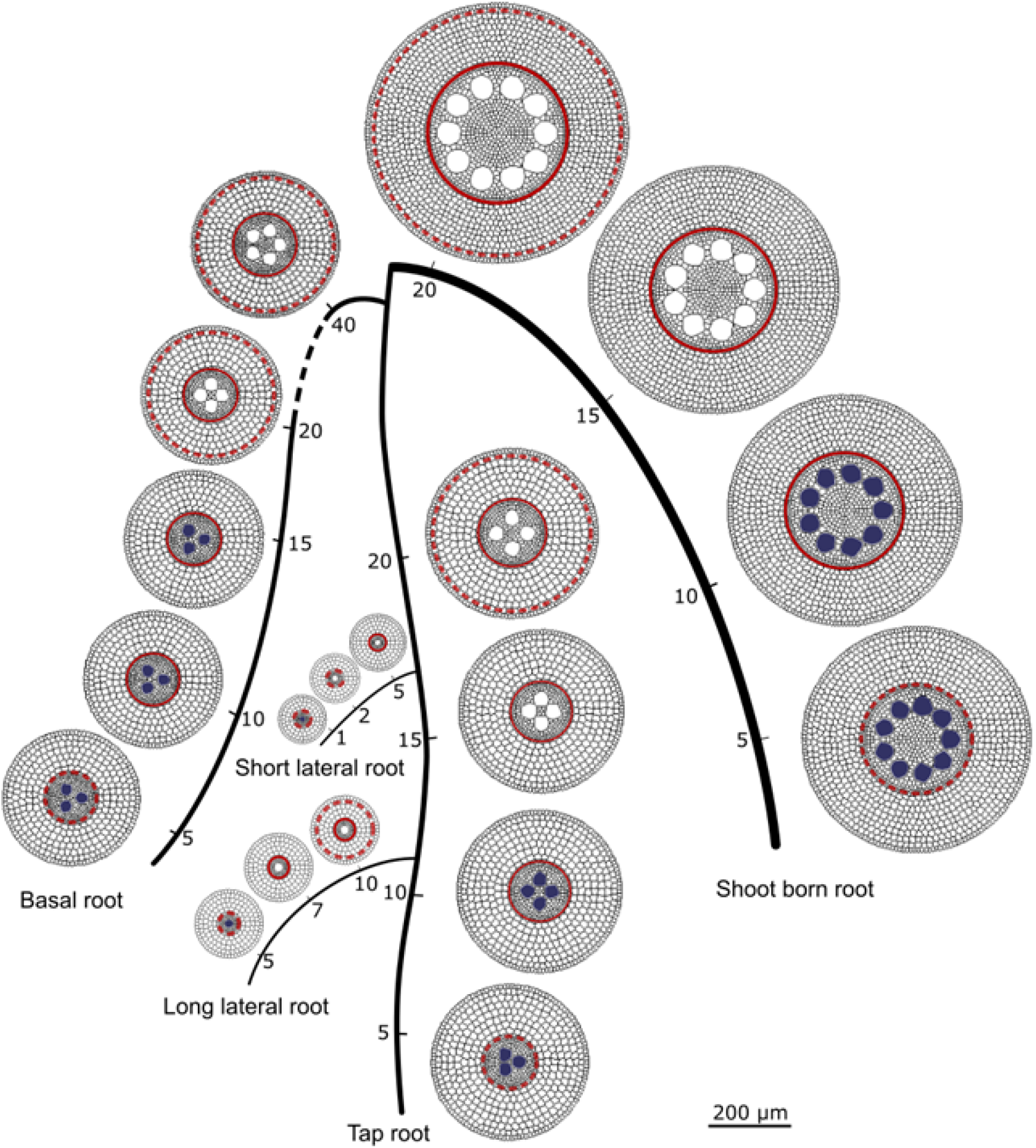
Schematic representation of a maize root system with five root types. Along each root type, the generated average root cross sections are placed accordingly. The number along the roots describe the distance from the tip of the root, the scale is free in between. The bar = 200μm is for root cross section representation. The filled metaxylem vessels represent the immature ones. The dashed red circles stand for the Casparian strip on le local root tissue. The continuous red circles stand for the fully suberize cell wall of the local root tissue.

In addition to the overview of the root cross section of the root system, we added the localisation of hydrophobic barriers and meta-xylem maturation zone based on staining signals (fig. 4). The berberine-aniline blue fluorescent staining procedure for suberin, lignin, and callose allowed us to estimate where the different maturation zones occur (fig. 5). On the main root axes, the tap-, the basal-, and the shoot born-root have a fully suberised endodermis before the maturation of the metaxylem. In addition, the lignification of the metaxylem vessels, usually occurred shortly after the complete suberisation of the endodermis. On the opposite, for lateral roots, the metaxylem vessels are lignified before the complet suberization of the endodermis. Moreover short lateral roots have a lignified metaxylem vessels before the suberin lamellae start to deposit on the cell walls of the endodermis. For long lateral roots, the lignified metaxylem vessels were found where some suberin lamellae start to deposit on the cell walls of the endodermis.

**Figure 4:**
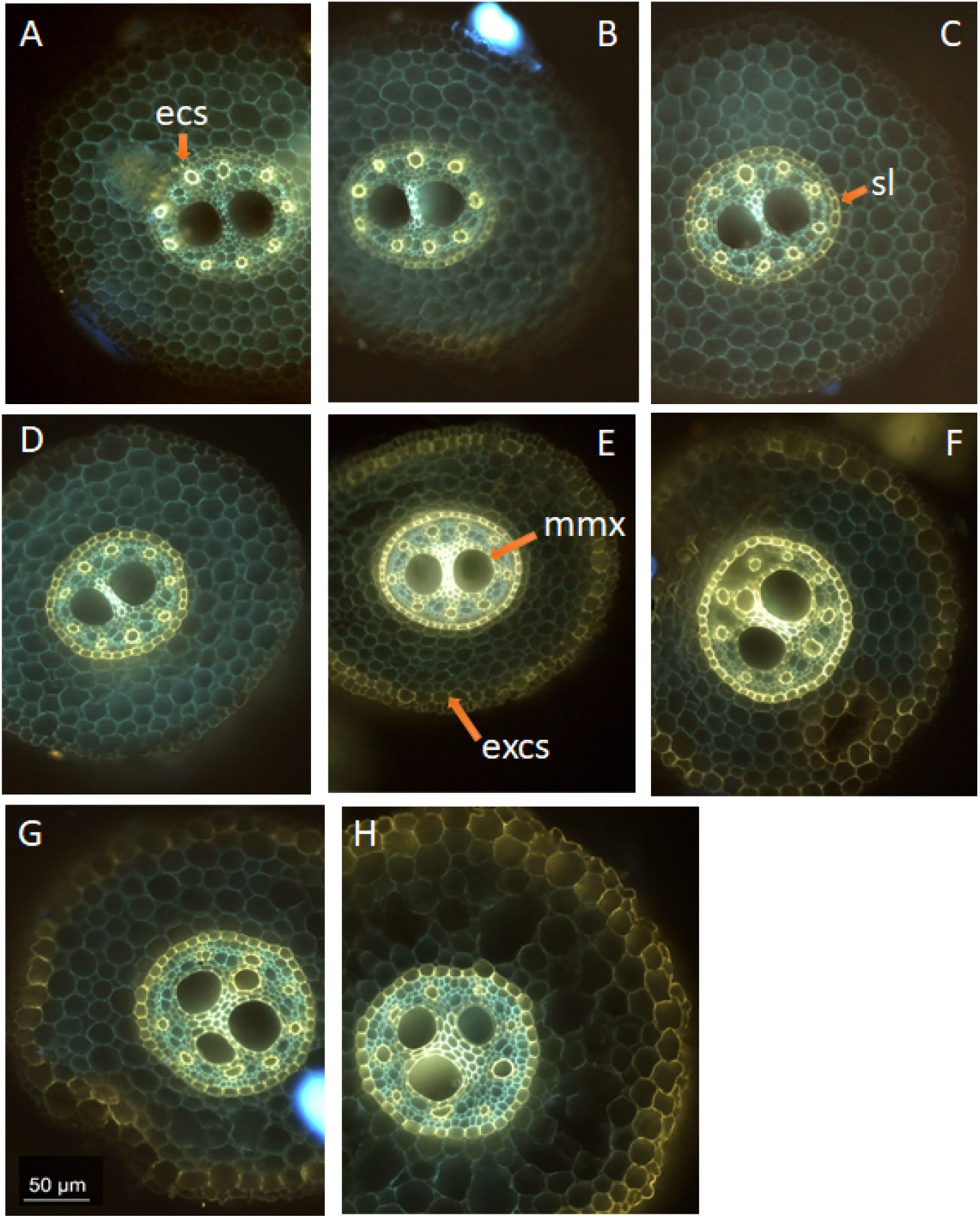
Basal root cross sections. A. 3 cm from apex, the arrow point at the endodermal Casparian strip (“ecs”); B. 5 cm from apex; C. 8 cm from apex, the arrow point at the suberin lamellae that formed on the endodermis (“sl”); D. 10 cm from apex; E. 15 cm from apex, the “mmx” arrow point at the lignify cell wall of the mature metaxylem vessels, the “excs” arrow point at the exodermal Casparian strip; F. 20 cm from apex; G. 25 cm from apex. H. 30 cm from apex. bar = 50 μm

**Figure 5:**
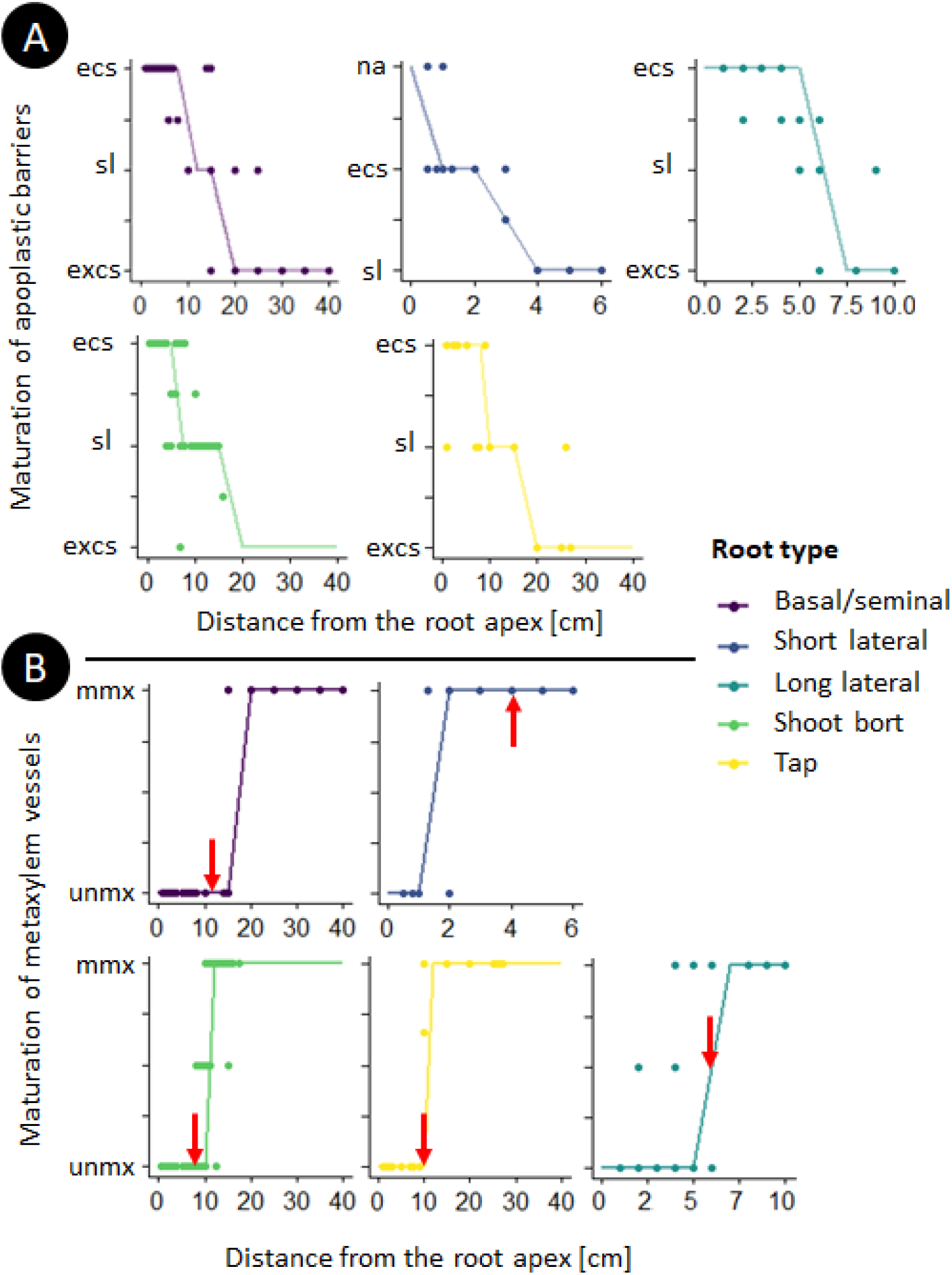
Evolution of the maturation for hydrophobic barriers (A) and for the metaxylem vessels (B) along the root axe for the different root types. Half values were applied when the transition between two maturations was observed. The lines are a discretization of the local weighted regressions of the scatter plots. (A) “na” = no hydrophobic barriers; “ecs” = endodermal Casparian strip; “sl” = fully suberized endodermis; “excs” = suberized endodermis and exodermal Casparian strip. (B) “unmx” = only the protoxylem vessels are lignified; “mmx” = All xylem vessels are lignified. The arrows point out where the endodermis is fully suberized for the specific root type.

### Hydraulic conductivity map

The next step of the process to make high resolution maps for the root hydraulic conductivity is to estimate the radial and axial conductivities of all the generated cross sections. To estimate the radial conductivity of the generated root cross section, we used the MECHA model (Couvreur et al., 2018) (fig. 6).

**Figure 6:**
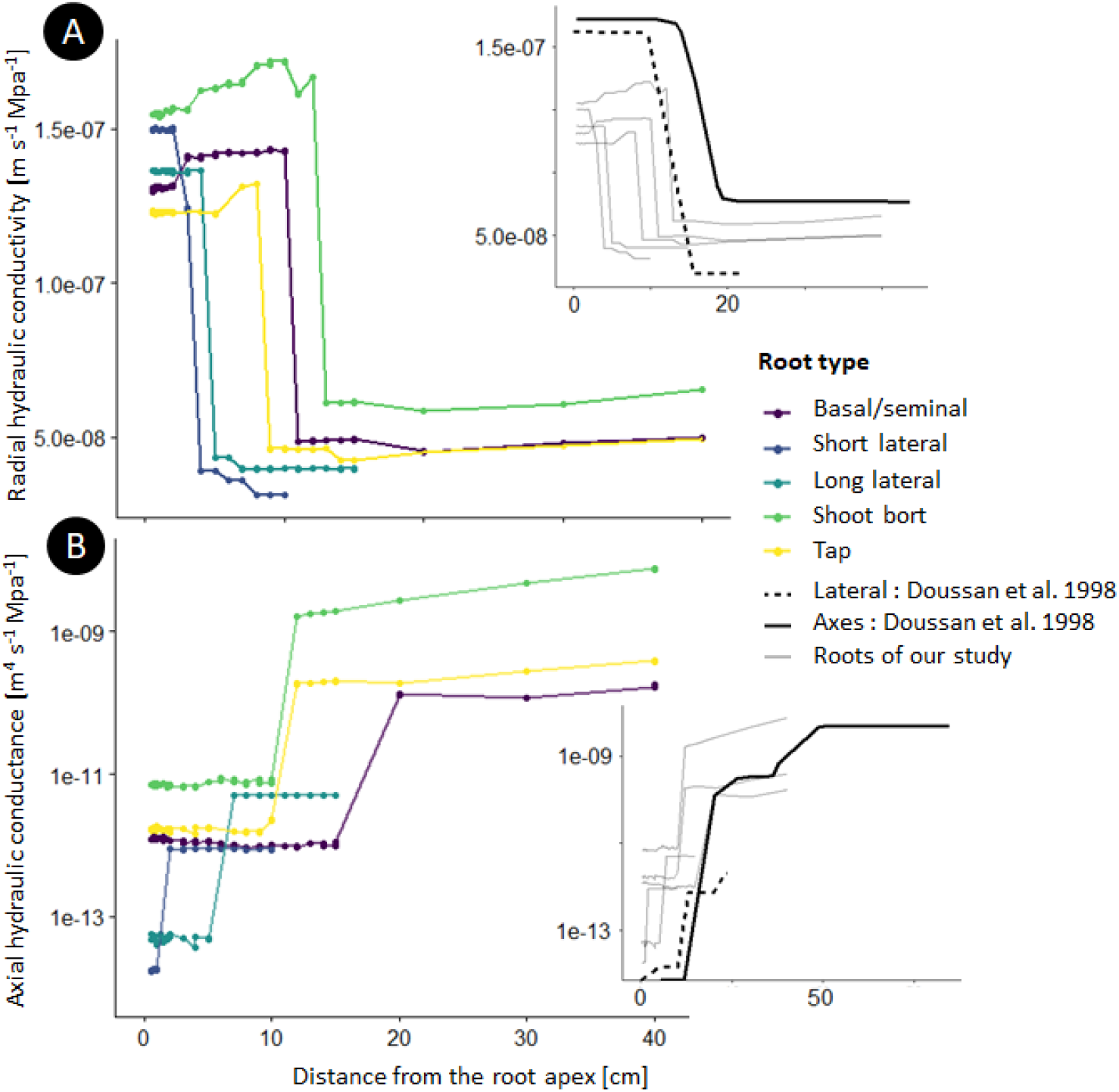
Hydraulic conductivity map for the different root types. A) Estimation of the radial hydraulic conductivity for each generated root cross section along the different root. The side graphic shows the two Doussan et al., 1998 estimations for *k_r_* and our estimations in comparison. B) Estimation of the axial hydraulic conductance for each generated root cross section along the different root. The side graphic shows the two Doussan et al., 1998 estimations for *k_x_* and our estimations in comparison.

We adjusted the maturation scenario in MECHA to fit our experimental data of the maturation zone for the hydrophobic barriers and metaxylem lignification. The cell hydraulic parameters were kept the same for all cross sections. For the axial conductivity, we used the Hagen-Poiseuille laws as explained in the Material and Methods section (equation 1 and 2).

## Discussion

In comparison to the hydraulic conductivity function of Doussan et al. (1998), our data, for the different root types, show a drop in radial conductivity closer to the tip. This is closer to the scenario of Zarebanadkouki et al. (2016), who estimated that the first drop occurred after four centimeters within the stepwise function with three transition zones. This early drop is due to the deposition of suberin lamellae in endodermal cell walls, which has been shown to be sensitive to environmental conditions (Tylová et al., 2017). The proportionally smaller second drop due to the addition of the exodermal Casparian strip is compensated further away by the expansion of the stele and the larger number of xylem vessels. Those anatomical effects on the radial conductivity follow the same trends as in Heymans et al. (2019). The radial conductivity estimations of our study are within range with the values of Doussan et al. (1998), and in a slightly higher range relative than the estimations by Zarebanadkouki et al. (2016) and Meunier et al. (2018).

The use of the Hagen-Poiseuille equations to estimate the *k_x_* is straightforward when the area of each xylem element is known. Our predicted range and trends both match direct measurements by Meunier et al. (2018) and estimations from Doussan et al. (1998). Uncertainties related to the application of the Hagen-Poiseuille law have been discussed in the literature. Frensch and Steudle (1989) have shown that it may overestimate experimental *k_x_* values by a factor of two to five. This could be due to the presence of perforation plates (Shane et al., 2000; Brodersen et al., 2018) or persistent xylem cross-walls (Sanderson et al., 1988). In this study we did not divide the estimated axial conductivity by a coefficient. The uncertainty of identification of mature xylem vessels by the used staining procedure could shift the transition zone shootward. We also assume that xylem sap has the same viscosity as water. This hypothesis could be discussed in relation to xylem sap temperature or solute concentration (Bruno and Sparapano, 2007).

The hydraulic conductivity map that we computed for this genotype in this precise environmental condition (*Zea mays var. B73* in pots) is an example case. Our methodology allows the inclusion of the effect of root anatomical changes and takes into account the selected cell hydraulic properties summarised in the material and methods section. The hydraulic conductivity map with five root types allows a better tuning for root water uptake models. This root hydraulic conductivity map can be used with other modelling tools to estimate other variables such as the root system conductance, or the standard sink fraction, as envisioned by Passot et al. (2019). Future inverse modelling studies could reuse the anatomical networks that we build on their root system architecture. Then, change in the modelling framework the cell hydraulic properties to match the macro hydraulic that would have been measured.

We developed a protocol that could be repeated in further studies (e.g. with different species, genotypes or environment). It is quicker than root pressure probing to estimate radial water flow. GRANAR takes around one to twenty seconds to generate root cross sections that are presented in this study. MECHA takes around one to five min per root cross sections to estimate the *k_r_*. On the opposite, one estimation for the *k_r_* from the root pressure probe takes at least three to five hours as steady root pressure has to be established after the connection between the root and the device (Liu et al., 2009). In both cases, making free-hand root cross-section takes around 10 to 20 minutes.

Meunier et al. (2020) showed that modifying hydraulic properties changes the root system hydraulic architecture and thus affects the whole root system conductance (*K_rs_*). Tuning root hydraulic conductivity functions to match experimental data or test new hypotheses through simulation studies could therefore show the local impact of root anatomy or cell hydraulic properties on the whole root conductance. A better understanding of the effect of local root traits on the global hydraulic behaviour of the root system could enhance the breeding efforts towards more drought tolerant cultivars.

## Conclusion

In this study, we showed how to use stained root cross section images and computational tools (organ scale models: GRANAR and MECHA) to create high resolution hydraulic maps of a maize root system (var. B73 in our example). Our hydraulic map includes hydraulic information (radial and axial properties) and anatomical data along 5 root types (tap, basal, shoot born, long laterals and short laterals).

Anatomical differences along the root axes and between root types seems to have an impact on the radial and axial water flow through the roots. The values and trends shown in this study are in the same range as the estimations that can be found in the literature.

Side by side with measures from root pressure probing, our method has the advantages of being quick and output high resolution results. We expect our methodology to be of great use for further root hydraulic studies. It will help match the hydraulic conductivities of root systems and experimental data, or test new hypotheses through simulation studies. These local root conductivities can be used in functional-structural root models to estimate macro hydraulic properties. It launches synthetic ways to test or benchmark the local impact of local root traits on the global hydraulic behaviour of a root system.

## Abbreviation

*k_AQP_*: contribution of aquaporins to the plasma membrane hydraulic conductivity
*k_m_*: plasma membrane intrinsic hydraulic conductivity
*K_PD_*: conductance of plasmodesmata per unit membrane surface
*k_r_*: radial hydraulic conductivity
*K_rs_*: root system hydraulic conductance
*k_x_*: specific axial hydraulic conductance
*L_pc_*: protoplast permeability
tip: root apex

